# The social evolution of siderophore production in *Pseudomonas aeruginosa* is environmentally determined

**DOI:** 10.1101/095323

**Authors:** Freya Harrison, Alan McNally, Ana C. da Silva, Stephen P. Diggle

## Abstract

Bacteria secrete various exoproducts whose benefits can be shared by all cells in the vicinity. The potential importance of these “public goods” in bacterial evolutionary ecology has been extensively studied. Cheating by siderophore-null mutants of the opportunistic pathogen *Pseudomonas aeruginosa* has received particular attention. The potential of siderophore mutants to attenuate virulence, and the possibility of exploiting this for clinical ends, have generated a wealth of publications. However, the possibility that genotype · environment interactions govern the evolutionary consequences of siderophore loss has been almost entirely ignored. A review of the available literature revealed (i) widespread use of an undefined mutant as a siderophore cheat; and (ii) a reliance on experiments conducted in iron-limited minimal medium. Whole genome sequencing of the undefined mutant revealed a range of mutations affecting phenotypes other than siderophore production. We then conducted cheating assays using defined deletion mutants, grown in conditions designed to model infected fluids and tissue in CF lung infection and non-healing wounds. Depending on the environment, we found that siderophore loss could lead to cheating, simple fitness defects, or no fitness effect at all. It is therefore crucial to develop appropriate *in vitro* growth conditions in order to better predict the social evolution of traits *in vivo*.

## Introduction

Bacteria are social organisms, displaying coordinated behaviours including quorum sensing (QS), biofilm formation, and the production of shareable exoproducts (West et al 2007). Exoproducts which act as “public goods” (including iron-scavenging siderophores, exoproteases, biofilm polymers, toxins and QS signals) are vulnerable to “cheating”: cells which do not produce a particular molecule avoid the costs of production but reap the benefits of their neighbours’ investment and increase their evolutionary fitness at the expense of producers (Darch et al 2012, Diggle et al 2007, Griffin et al 2004, Harrison et al 2006, Jiricny et al 2010, Mund et al in prep, Raymond et al 2012). The circumstances under which cooperation can be maintained in the population over evolutionary time, versus those under which cheating prevails, have been predicted by theory (Frank 1998, Hamilton 1964). Because bacteria are amenable to evolution experiments, they have been widely used to test theoretical predictions (e.g. Diggle et al 2007, Griffin et al 2004, Kümmerli et al 2009a). Siderophore production by the opportunistic pathogen *Pseudomonas aeruginosa* is a particularly tractable workhorse for sociomicrobiology, facilitating tests of key evolutionary hypotheses (Table 1).

**Table 1.**
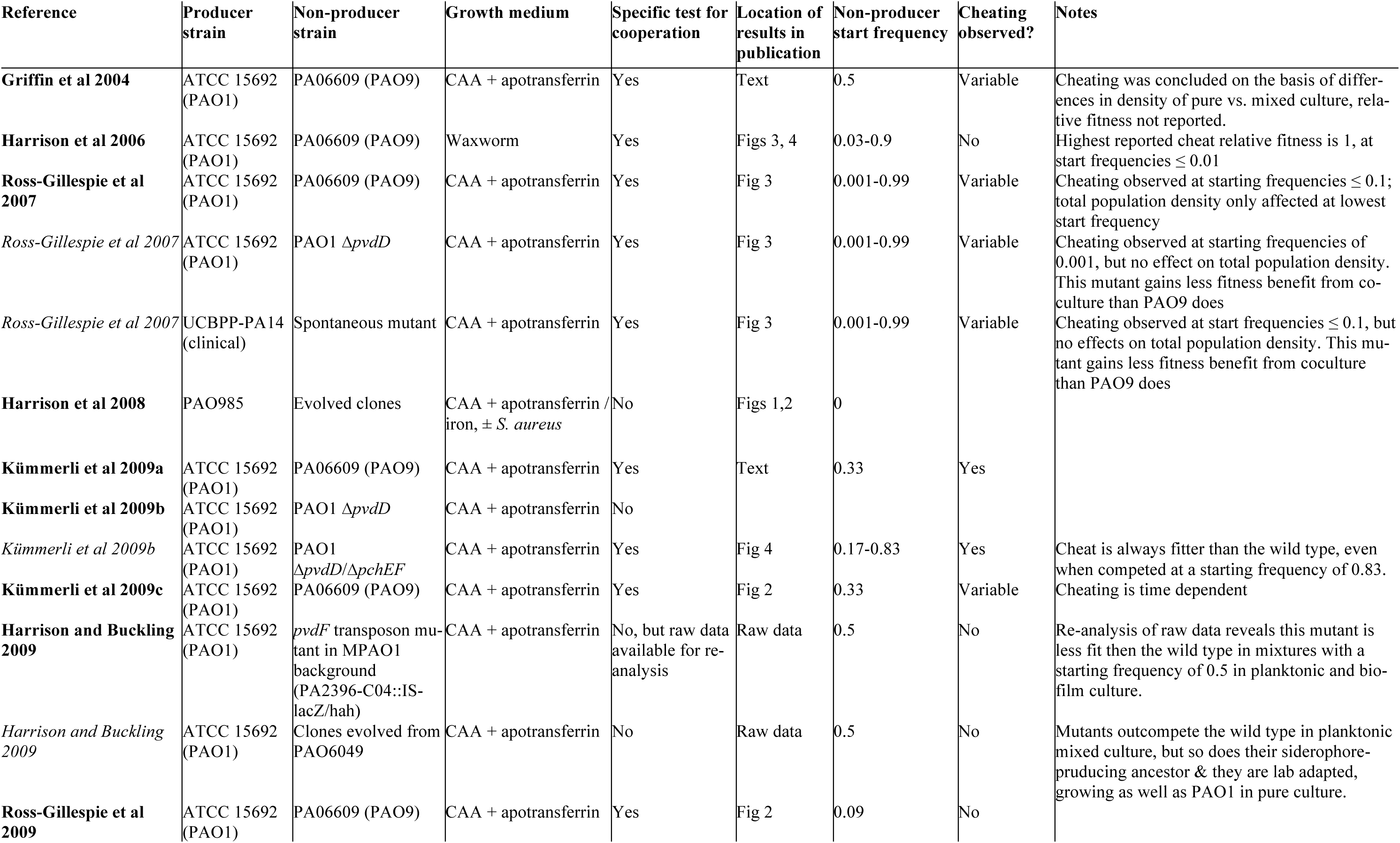
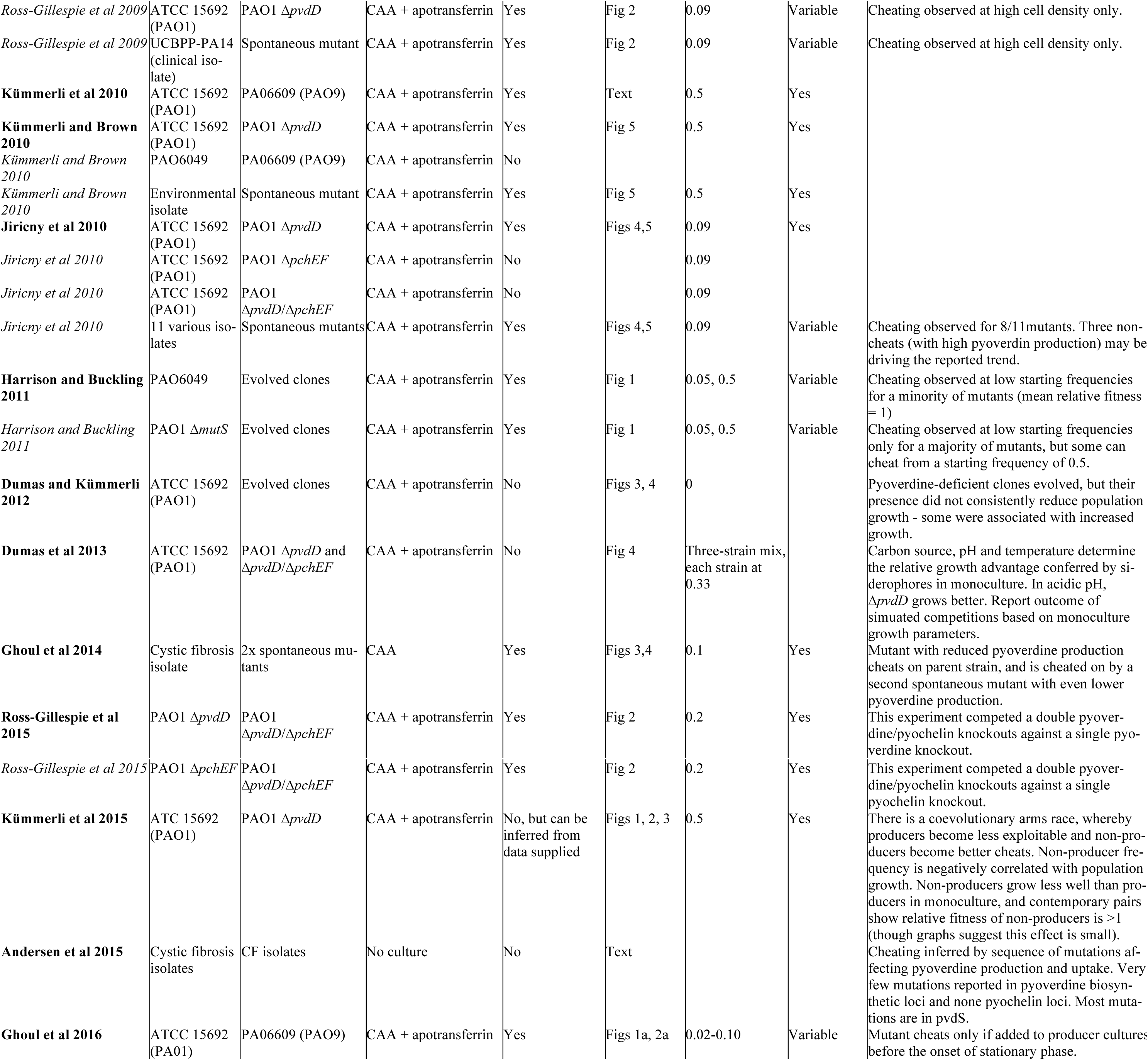
Results of a review of the empirical literature on *P. aeruginosa* siderophore mutants and cheating.

Early research into bacterial public goods focussed on using them as models to test general theory. But as the field progressed, researchers began to suggest that public goods play important roles in infected hosts. This led to suggestions that cooperation among pathogens could be manipulated for clinical ends, especially in hard-to-treat chronic infections (e.g. Harrison et al 2006, Rumbaugh et al 2009, Brown et al 2009, Foster 2005). *P. aeruginosa* causes various chronic infections in immunocompromised hosts, notably lung infections in people with cystic fibrosis (CF) and wound infections in people with burns or diabetic ulcers (Friedrich et al 2015, Hirsch and Tam 2010). Siderophore-null mutants can invade *in vitro* populations and cheat their way to high density, resulting in population reduction. Researchers have hypothesised that the presence of siderophore mutants in chronic infection isolates can be explained by cheating *in vivo*, and that siderophore mutants could be used as “Trojan horses” to ferry antibiotic-susceptibility alleles into infections and thus render them curable (Brown et al 2009, Harrison et al 2006).

One potential problem with this reasoning is the implicit assumption that gene expression and the fitness consequences of mutations are comparable *in vitro* and *in vivo.* Technical advances have revealed just how much bacterial transcriptomes (Croucher and Thomson 2010) and the fitness of loss-of-function mutants (Van Opijnen et al 2009) vary across environments; indeed, environment may be a bigger predictor of *P. aeruginosa* phenotype than its genetic background (Dötsch et al 2015). When the environment bacteria experience in a specific infection context is carefully modelled in the lab, the importance of genotype × environment interactions in determining fitness become very clear (Harrison et al 2014, Palmer et al 2007, Turner et al 2015).

Typically, siderophore experiments are conducted in iron-limited minimal medium, creating clear costs and benefits to siderophore production. While this may recapitulate the iron restriction encountered by pathogens colonising healthy hosts during acute infection, chronic infections present an entirely different environment. Tissue damage, a hyper-inflammatory response and disease-specific changes in host phenotype (e.g. increased mucus volume and adhesivity in CF lungs, (Boucher 2007) are accompanied changes in the growth substrates available to bacteria. In CF lungs, bacteria use amino acids released by damaged tissues, or from mucus, as carbon sources (Flynn et al 2016, Palmer et al 2005, Palmer et al 2007), and iron is plentiful (Tyrrell and Callaghan 2016). Consequently, bacterial gene expression – and presumably the roles played by virulence factors – differ in chronic and acute contexts (Goodman et al 2004). Because lab models of the environments encountered in CF lungs and soft-tissue wounds have been developed (Harrison et al 2014, Harrison and Diggle in press, Palmer et al 2007, Turner et al 2015, Werthén et al 2010), there is a pressing need to re-assess the role played by siderophore mutants in environments that better model chronic infection.

To assess and extend the potential of laboratory experiments to yield clinically useful data on the evolutionary dynamics of siderophore production, we defined three aims. First, we reviewed published experimental work on *P. aeruginosa* siderophore cooperation. This literature has not previously been systematically reviewed. Our goal was to characterise any biases in the literature which could restrict its applicability to the various environments in which this flexible opportunist can thrive. We found two such biases and defined two further empirical aims to address these.

The first potential bias in the literature was that many experiments (by ourselves and others) used an uncharacterised, UV-generated mutant (PAO9) as a siderophore cheat. We therefore conducted whole-genome sequencing of this mutant to determine (i) the genetic basis of its siderophore-null phenotype, and (ii) whether it carries other mutations that could affect the outcome of competition with the wild type. The second source of bias was the lack of studies employing a well-defined model of chronic infection. We therefore determined the fitness consequences of siderophore loss-of-function mutations in relatively well-characterised models of CF lung infection and chronic soft-tissue wounds. Because we found many mutations that potentially affect metabolism and growth in PAO9, we used siderophore deletion mutant of the siderophores pyoverdine and pyochelin for these experiments. CF lung infection was modelled using (a) liquid artificial sputum medium (ASM: Palmer et al 2007, Turner et al 2015); and (b) an *ex vivo* model of biofilm infection in which small sections of porcine bronchiole are infected with bacteria and cultured in ASM (Harrison and Diggle in press). Chronic wound infection was modelled using (a) synthetic wound fluid (SWF), and (b) synthetic wound fluid solidified with collagen to form model soft-tissue plugs (Werthén et al 2010). We explicitly explored differences between liquid medium and structured infection models because spatial structure affects the social dynamics of public goods (Frank 1998, Kümmerli et al 2009c, Mund et al in prep), and because biofilm formation in structured models could alter gene expression and the fitness consequences of mutation (Bjarnsholt et al 2013, Whiteley et al 2001). Our results demonstrate that whether siderophore mutants act as cheats is dependent on the environment, and suggest that these mutants may not act as cheats *in vivo.*

## Meaterials & Methods

### Bacterial strains

*P. aeruginosa* ATCC 15692 (PAO1) was used as a wild type siderophore producer and clean Δ*pvdD* and Δ*pvdD*Δ*pchEF* mutants in this background (Ghysels et al 2004) used as single pyoverdine and double pyoverdine/pyochelin mutants, respectively. The UV-induced mutant PA6609 (PAO9) was derived from PAO6049 (Hohnadel et al 1986); PAO6049 is a Tn5-induced methionine auxotroph derived from PAO1 (Rella et al 1985).

### Collating and analysing experimental work on siderophore cheating

To identify published experiments on siderophore cheating, we used Scopus.com to (i) conduct a literature search and (ii) locate articles citing already-identified experimental and theoretical articles on siderophore social evolution. For each experiment in each article, we recorded which strains/clones and media were used; whether an explicit test for cooperation was conducted (relative fitness of mutant in pure/mixed culture with a producer strain & growth or final density of pure/mixed cultures); the starting frequency of mutants in coculture experiments; the location of key data in the article; whether cheating was observed and any other key conclusions. The diverse starting frequencies and mutants used make this data unsuitable for formal meta-analysis, so we simply present the key characteristics and findings of the studies in a format that facilitates qualitative comparison.

### Whole genome sequencing of PAO6609 (PAO9)

PAO9 was cultured overnight in 10 ml Lysogeny Broth at 37°C on an orbital shaker. Genomic DNA was extracted using a Sigma Aldrich GenElute Bacterial Genomic DNA Kit. Library preparation was performed using the Nextera XT library preparation kit (Illumina), and 2 x 300 bp paired-end sequencing performed on the Illumina MiSeq platform using a V3 sequencing cartridge, to approximately 50x coverage. Reads were assembled using SPAdes run with the –careful flag, and annotated using Prokka. This produced an assembly of 6,232,039 bp comprising 89 contigs with an N50 of 220,789. The assembly was compared with the reference PAO1 genome (NCBI reference sequence NC_002516) using BLAST and the comparison visualised using ACT to search for gene acquisition or loss events. SNP typing was performed by mapping the raw reads of PAO9 against the PAO1 reference genome (NC_002516.2) using SMALT and Samtools. A total of 98.3% of 1.17M reads mapped to the reference, from which high fidelity SNPs were called using a cut off of minimum allele frequency of 0.8, minimum quality score 30, and minimum depth of 8.

### Growth conditions for cheating experiments

We used five different growth environments. In each case, cultures were grown in 24-well microtitre plates and incubated on a rocking platform at 37°C. (i) 2 ml casamino acids medium (CAA: 5 g casamino acids, 1.18g K_2_HP0_4_·3H_2_0, 0.25 g MgSO_4_·7H_2_O, per litre) supplemented with 20 mM NaHCO_3_ and 100 μg mL^-1^ human apo-transferrin (Sigma), cultured for 24 hours. (ii) 2 ml artificial sputum medium (ASM) following recipe in Palmer et al (2007), cultured for 24 or 48 hours. (iii) 5 mm^2^ pig bronchiole + 400 μ1 ASM following protocol in Harrison and Diggle (in press), cultured for 96 hours. (iv) 2 ml synthetic wound fluid (50% v/v peptone water / fetal bovine serum following Werthén et al (2010)), cultured for 24 or 48 hours. (v) 400 μ1 synthetic chronic wound (SCW: SWF solidified with rat tail collagen following protocol in Werthén et al, (2010)), cultured for 48 hours. For environments ii-v, we measured wild-type siderophore production and calculated the fitness of siderophore mutants relative to the wild type in clonal and mixed culture. Four or five replica cultures were inoculated with (a) wild-type, (b) Δ*pvdD* (c) Δ*pvdD*Δ*pchEF*, (d) 50% wild-type + 50% Δ*pvdD* or (e) 50% wild-type + 50% Δ*pvdD*Δ*pchEF* bacteria for each environment/culture time combination, and each experiments was repeated twice to yield two experimental blocks. For environment i (CAA), we simply measured siderophore production by four replica populations of the wild type for comparison with environments ii-v.

The density of the inoculum was kept as consistent as possible across environments and population types, as cell density can affect the outcome of wild-type – siderophore-null co-culture experiments (Ross-Gillespie et al 2009). For experiments in CAA, ASM and SWF, each culture was inoculated with a total of two colonies of the relevant genotype(s), picked from LB plates using a 200 μl pipette tip. For experiments in *ex vivo* bronchiole and SCW, each culture was inoculated with two colonies of the relevant genotype(s), picked from LB plates using an insulin syringe fitted with a 30 G needle.

To construct time courses of bacterial growth in ASM, two colonies of (a) wild-type, (b) Δ*pvdD* or (c) Δ*pvdD*Δ*pchEF* were inoculated into four 2 ml aliquots of ASM in a 24-well microtitre plate. This was incubated at 37°C in a Tecan Infinite 200 Pro multimode reader for 18 hours; every 20 minutes, the plate was shaken for two seconds and absorbance of each well read at 400 nm. (Reading at more traditional values of 600-700 nm risks interference from absorbance by pyoverdine in wild-type cultures).

### Statistical analyses

Data were analysed in R 3.3.0 (Team 2016) using general linear models and ANOVA. To meet model assumptions of homoscedasticity and normality of residuals, fitness data from SWF were log-transformed and fitness data from ASM and *ex vivo* bronchiole square-root transformed; fitness data from SCWs did not require transformation. Where missing values caused non-orthogonality, the *car* package (Fox and Weisberg 2011) was used to perform ANOVA with Type II sums of squares, returning correct F-ratios for main effects in models containing an interaction. Fitted means and confidence intervals were retrieved, and post-hoc contrasts tested, using the *lsmeans* package (Lenth 2013). Post-hoc multiple comparisons against a control were implemented using Dunnett’s test in the *multcomp* package (Hothorn et al 2008).

### Data availability

Raw sequence data has been deposited in the European Nucleotide Archive (Accession number: ERR1725797). Summary information on mutations in PAO9 (Fig 1) and raw data for experiments depicted in Figures 2-5 and S2 are available as Supplementary Information at the *Nature Ecology & Evolution* website.

**Figure 1.**
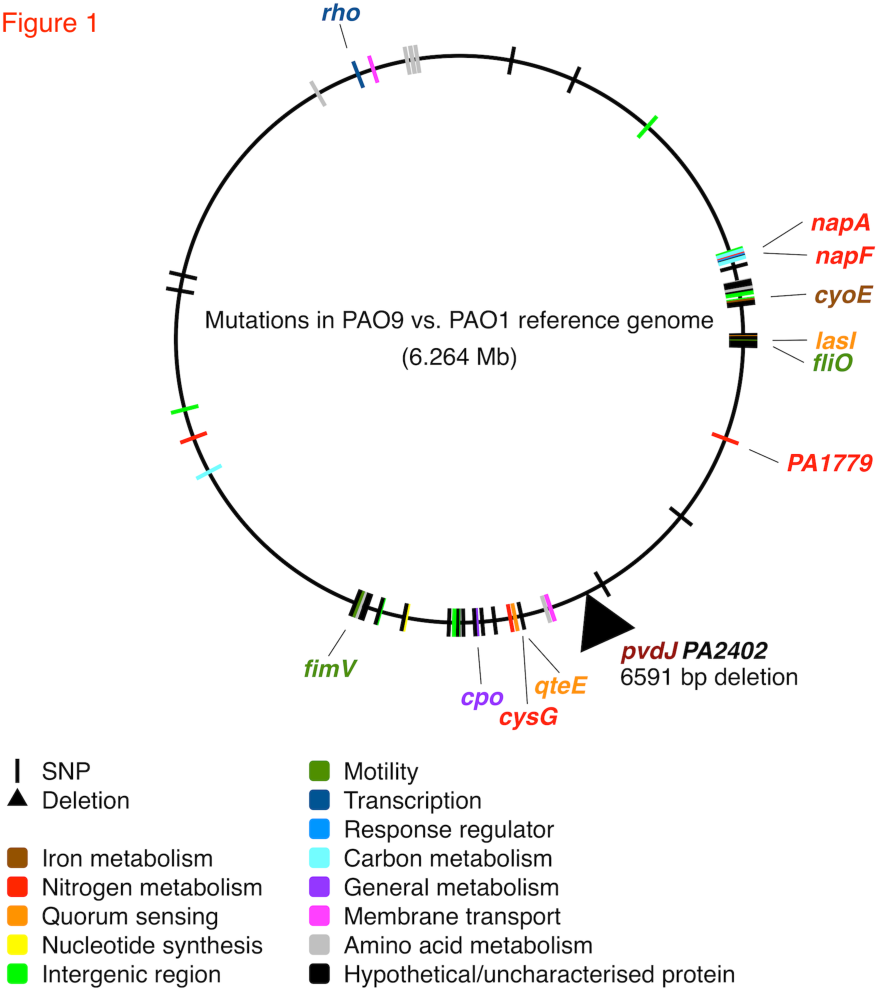
Location of one deletion and 90 SNPs in the genome of PAO6609 (PAO9), mapped against a PAO1 reference sequence (NC_002516.2). SNPs are colour coded by functional class of the locus affected. 11 SNPs that that result in amino acid substitution and which seem most likely to affect growth and virulence are highlighted and the gene name given.

**Figure 2.**
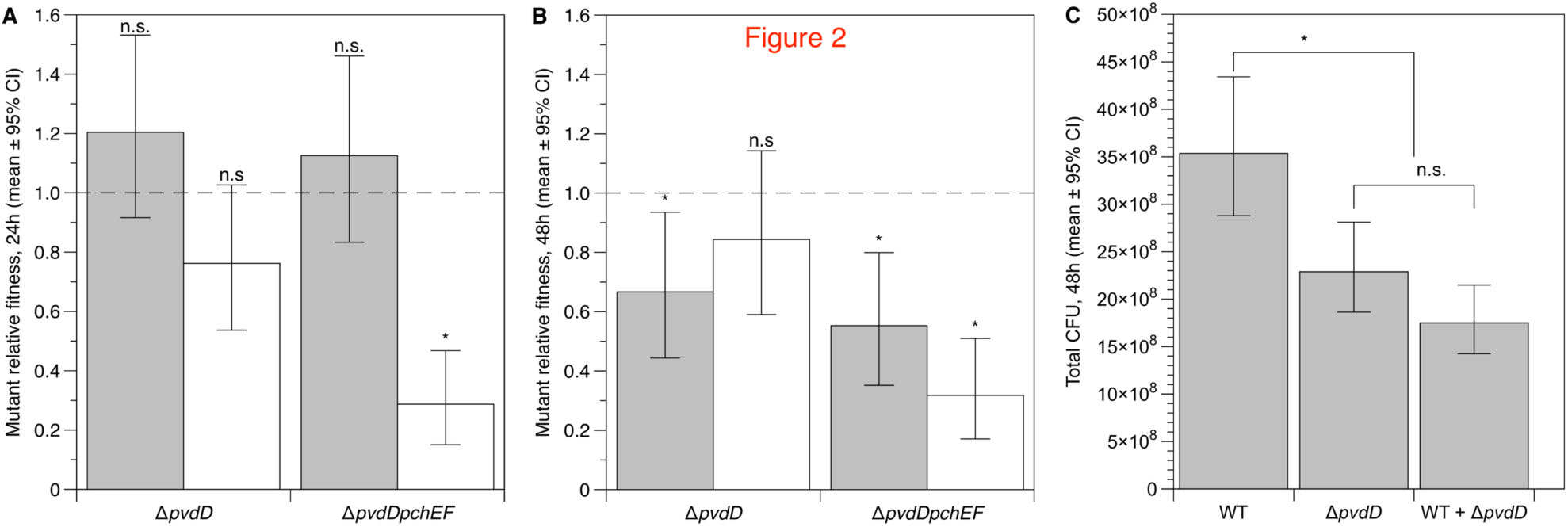
(a,b) Relative fitness of Δ*pvdD* and *ΔpvdD ΔpchEF* mutants in pure culture (grey bars) and in mixed culture with an isogenic wild type (white bars) in ASM after (a) 24 and (b) 48 hours of growth. (c) Cheating by Δ*pvdD* over 48 hours of co-culture results in mixed wild type + mutant cultures showing the same reduction in total population density as pure mutant cultures. Bars show means of 9-10 replicates split across two replica experiments, with associated 95% confidence interval. After ANOVA, post-hoc tests were conducted to determine whether each fitted mean value was significantly different from 1: * = *p* < 0.02; n.s. = not significant.

**Figure 3.**
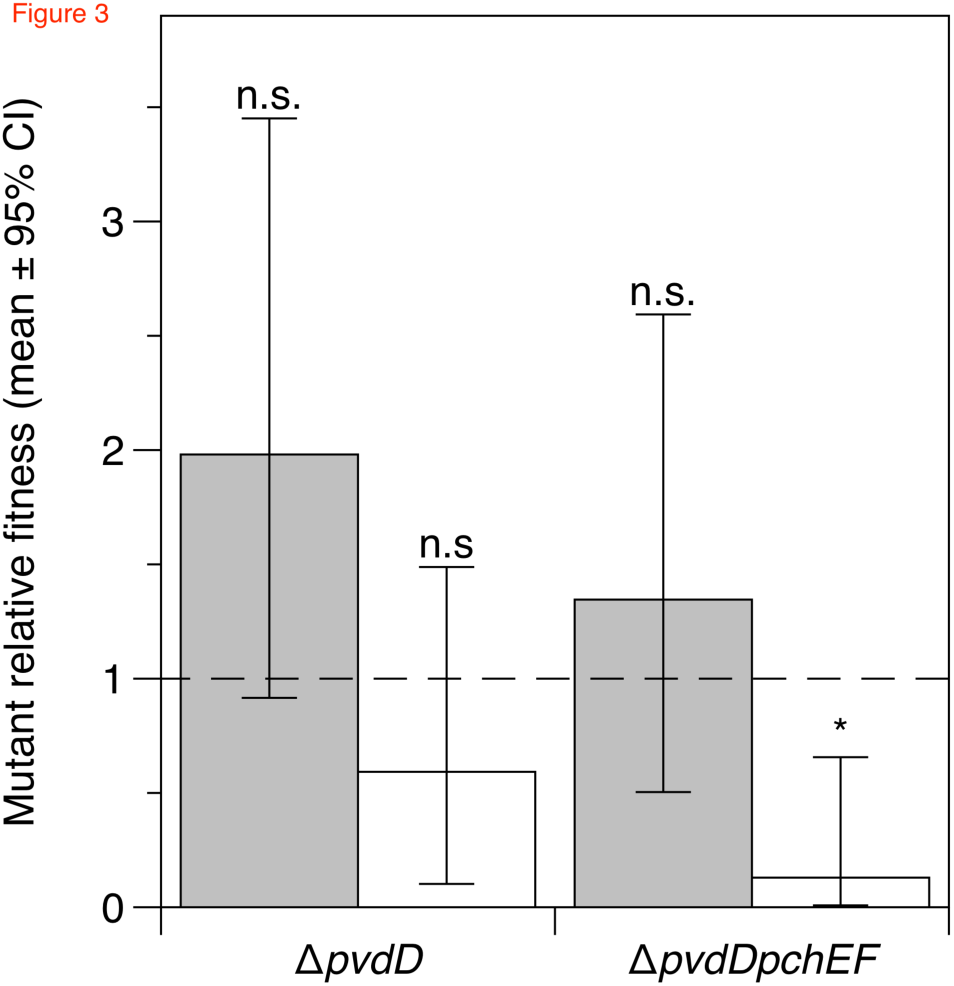
Relative fitness of Δ*pvdD* and Δ*pvdD*Δ*pchEF* mutants in pure culture (grey bars) and in mixed culture with an isogenic wild type (white bars) in *ex vivo* pig lung + ASM after 96 hours of growth. Bars show means of 8 replicates spread across two replica experiments, with associated 95% confidence interval. After ANOVA, post-hoc *t*-tests were conducted to determine whether each fitted mean value was significantly different from 1: * = *p* = 0.007; n.s. = not significant.

**Figure 4.**
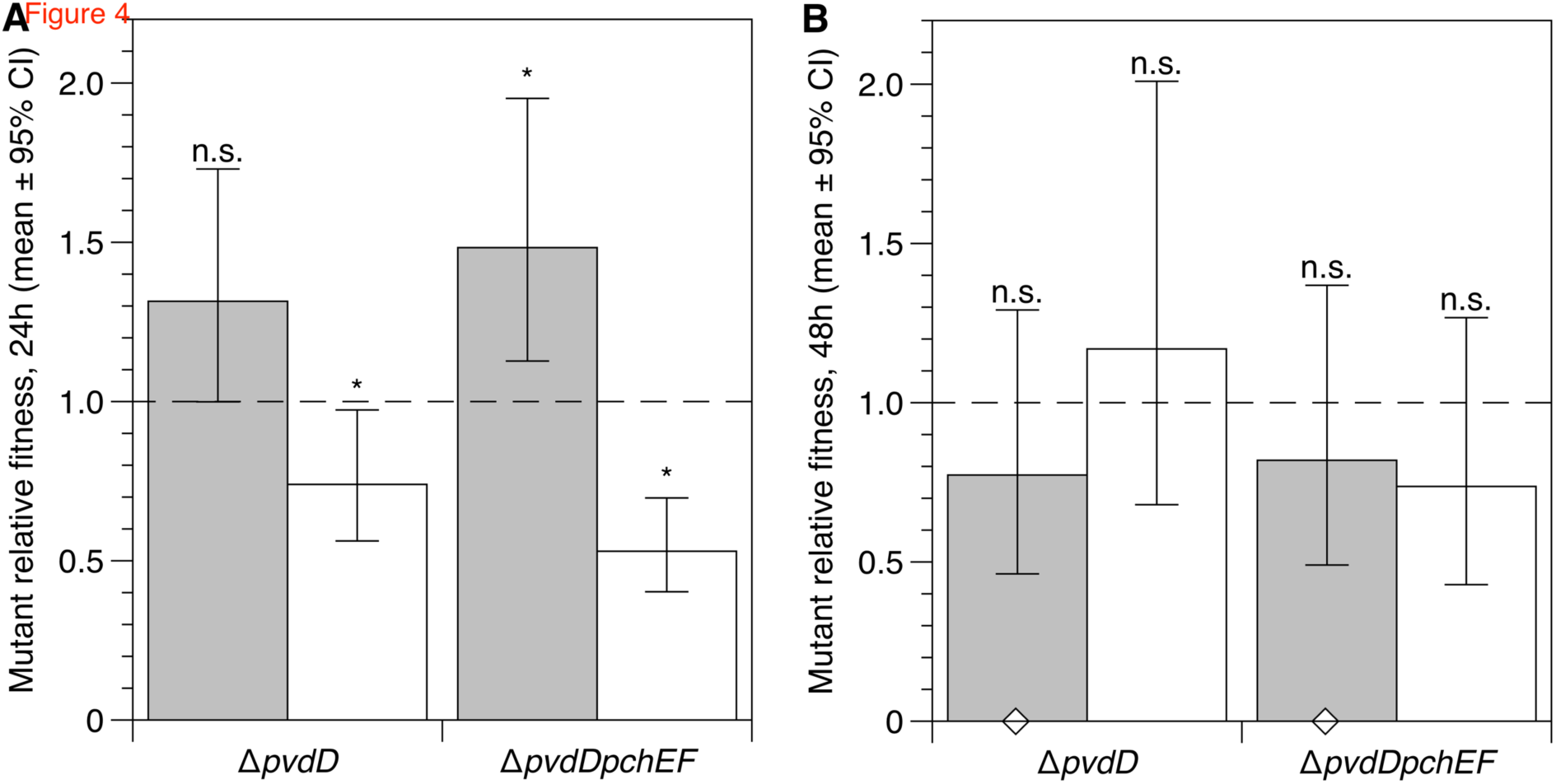
Relative fitness of Δ*pvdD* and Δ*pvdD*Δ*pchEF* mutants in pure culture (grey bars) and in mixed culture with an isogenic wild type (white bars) in SWF after (a) 24 and (b) 48 hours of growth. Bars show means of 10 replicates split across two replica experiments, with associated 95% confidence interval. After ANOVA, post-hoc *t*-tests were conducted to determine whether each fitted mean value was significantly different from 1: * = *p* < 0.02; n.s. = not significant. Confidence intervals and *p*-values in (b) are taken from a model that excluded two outliers where relative fitness was zero (indicated by open diamonds): including these outliers in the analysis did not change any of the conclusions given in the text.

**Figure 5.**
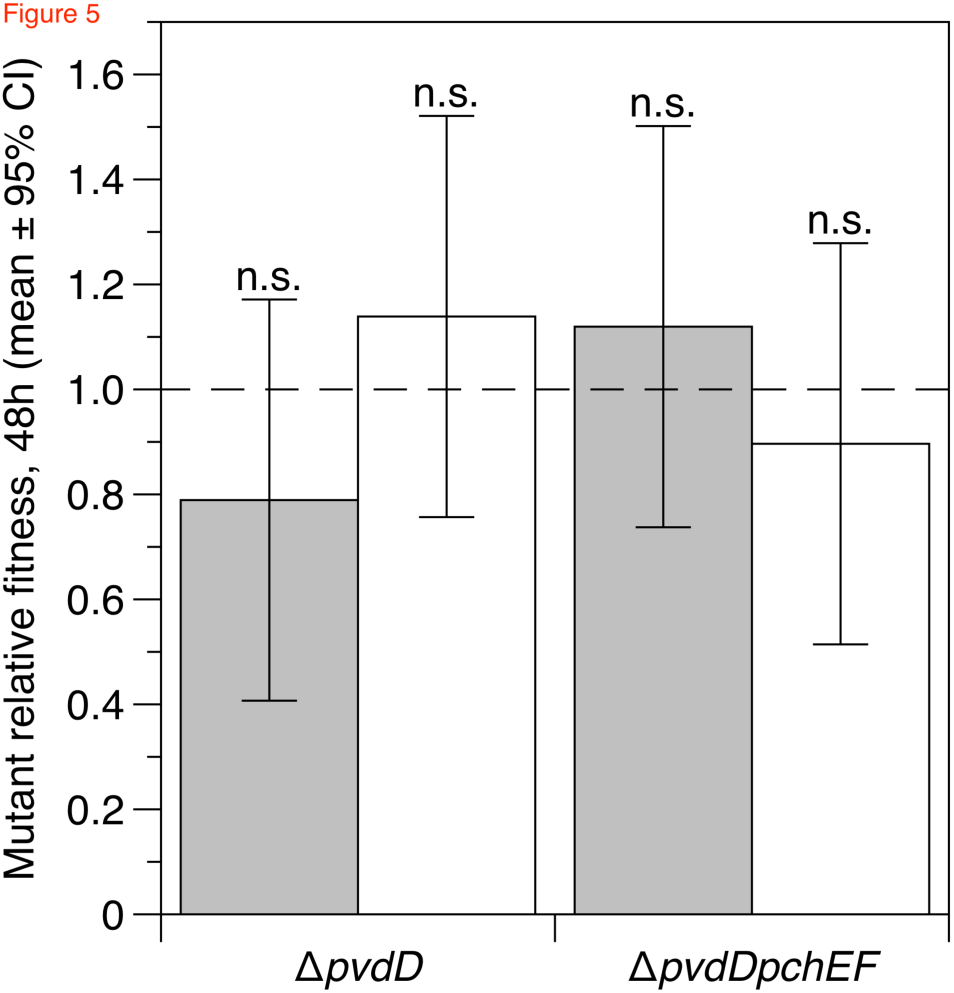
Relative fitness of Δ*pvdD* and Δ*pvdD*Δ*pchEF* mutants in pure culture (grey bars) and in mixed culture with an isogenic wild type (white bars) in synthetic wounds after 48 hours of growth. Bars show means of 10 replicates split across two replica experiments, with associated 95% confidence interval. After ANOVA, post-hoc *t*-tests were conducted to determine whether each fitted mean value was significantly different from 1: n.s. = not significant.

## Results

### Defining key research priorities: analysis of published experiments

A review of published experiments on *P. aeruginosa* siderophore mutants and cheating identified 33 experiments in 18 published articles, summarised in Table 1. Two-thirds of the experiments used the well-characterised lab strain PAO1 as the siderophore-producing wild-type, and 5 experiments used clinical or environmental isolates. Of the 28 experiments that did not focus on natural isolates, 14 used defined siderophore mutants. These included deletions of the *pvdD/pvdF* and/or *pchEF* loci, which are involved in the biosynthesis of the siderophores pyoverdine and the pyochelin, respectively. Of the remainder, 9 used the UV-induced mutant PAO9 and 5 used spontaneous or evolved mutants.

31/33 experiments in Table 1 were conducted in CAA minimal medium, in all but one case made further iron-limited by the addition of the human iron chelator transferrin. This medium was not designed to reflect any specific natural environment: it was optimised to ensure that siderophores were necessary for growth, and acted as a public good, when bacteria were cultured in it (Griffin et al 2004). This makes it ideal for experiments to test evolutionary theory about why and how different social strategies can be selected for. Experiments in iron-limited CAA have, for instance, revealed that the potential for siderophore mutants to cheat can be curtailed by the scale of competition in a metapopulation (Griffin et al 2004) or by growth in a structured environment, where spatial segregation prevents mutants from accessing siderophores produced by wild types (Kümmerli et al, 2009c). As an opportunist with a large, flexible metabolome and secretome, *P. aeruginosa* can persist in a range of environments. Some of these may be approximated by iron-limited minimal media. For instance, experiments conducted in CAA may therefore be useful for understanding the dynamics of siderophore genotypes/phenotypes at the onset of acute infection in healthy host tissues, where iron is sequestered by high-affinity host chelators (including transferrin), or in nutrient-poor abiotic environments.

One study explicitly tested the effect of environmental variables on siderophore production in monoculture. Dumas et al (2013) dissected the relative roles of pyoverdine and pyochelin, suggesting that while pyoverdine is the primary siderophore expressed under severe iron limitation, under moderate iron limitation pyochelin production dominates. The authors used parameters estimated from these experiments to simulate wild-type – siderophore mutant competition in a range of environmental scenarios, and found that the fitness consequences of siderophore-null mutations varied. Based on these results, the authors suggested that pyoverdine-deficient mutants may be favoured in chronic infection due to better growth in high-iron, low-pH environments, rather than because they are cheats. The importance of environmental iron regime and pyoverdine/pyochelin switching in determining the fitness consequences of siderophore mutants has also been explored in a suite of experiments using the closely-related bacterium *P. fluorescens*: pyoverdine mutants of this species were shown to cheat under severe, but not moderate, iron limitation (Zhang and Rainey 2013, Kümmerli and Ross-Gillespie 2013).

Further, the strength of siderophore cheating may be constrained. Cheating could become self-limiting as increases in mutant frequency alter the cost:benefit ratio of losing siderophore production, or the cost:benefit ratio may be different in actively-growing versus established populations. In many studies that explored a range of starting frequencies, mutants acts cheats only when they are initially inoculated at frequencies ≤ 0.1, which raises questions about the ability of these mutants to retain cheat at, and persist at, higher frequencies. In some cases, even though co-culture increased mutant relative fitness to > 1, there was no detrimental effect on total population density. These observations call into question the ability of siderophore mutants to act as “Trojan horses” to treat infection, as does the observation in one study (Ghoul et al 2016) that a siderophore mutant only cheats if added to wild-type cultures before stationary phase.

Therefore, whether siderophore-null mutants act as cheats in chronic infection remains an open question. It is possible that increased availability of iron in chronically-damaged tissues means that the benefits of siderophore production are reduced and that production is downregulated, removing any advantage to siderophore-null genotypes. Alternatively, siderophores may be beneficial but not exploitable by potential cheats: this could result from spatial structuring, persistence in stationary phase, and/or a reliance on siderophore recycling rather than continued production (Kümmerli & Brown 2010).

### Whole genome sequence of a commonly-used siderophore cheat, PAO9

The nature of the mutation leading to the siderophore-negative phenotype in PAO9 is not known. Further, as UV mutagenesis is non-specific, it is likely that this strain carries additional mutations that influence other important aspects of its phenotype, or moderate the fitness consequences siderophore loss. Without knowing the exact genotype of PAO9, we cannot be sure that empirical results obtained using it are the result of its siderophore phenotype alone. We performed whole genome sequencing of this strain and mapped the raw sequence data against a PAO1 reference genome (NCBI Reference Sequence: NC_002516.2, Stover et al 2000). The results are summarised in Figure 1. PAO9 harbours a single ˜6.6kb deletion, removing most of the non-ribosomal peptide synthetase locus *pvdJ*, which is involved in synthesising the pyoverdine side chain. This deletion also removes part of the open reading frame immediately upstream of *pvdJ*, PA2402. PA2402 encodes a probable non-ribosomal peptide synthetase of unknown function. We also found 90 high-resolution SNPs. The details of the deletion and SNPS are provided as Supplementary Information. Briefly, none of the SNPs were in loci associated with pyoverdine or pyochelin biosynthesis. However, we found SNPs resulting in mis-sense mutations in *cyoE*, which is involved in iron metabolism, in four loci associated with nitrogen metabolism (*napA*, *napF*, *cysG*, PA1779), two loci involved in quorum sensing *(lasI*, *qteE*), two loci involved in motility (*fliO*, *fimV*) and the *cpo* gene, which is involved in the biosynthesis of many organochlorine compounds: all of these mutations could potentially affect growth and/or virulence. Finally, we found a missense mutation in the *rho* transcription termination factor. Further work would be needed to deduce the effects of this mutation, but the possibility of generalised defects in the control of gene expression cannot be ruled out. In summary, PAO9 harbours numerous mutations in addition to the *pvdJ* partial deletion that likely make it an unreliable strain to use for empirical studies of the fitness and virulence consequences of siderophore loss.

### Production of siderophores by *P. aeruginosa* in models of chronic lung and wound infections

Because of the problems identified with PAO9, we used defined siderophore mutants to test the potential for social cheating in laboratory conditions that have been developed to represent specific chronic infection contexts, with strict attention to maximising likely ecological validity. As discussed above, while most research has focused on pyoverdine, under less severe iron limitation the metabolically cheaper siderophore pyochelin may take on a primary role in iron chelation (Dumas et al 2013, Kümmerli and Ross-Gillespie 2013). We therefore used a standard wild-type lab strain (PAO1) and two isogenic deletion mutants: a single pyoverdine knock-out (Δ*pvdD*) and a double pyoverdine/pyochelin knockout (Δ*pvdD*Δ*pchEF*) constructed using allelic exchange (Ghysels et al 2004). Both mutants have been reported to act as social cheats in iron-limited CAA, when inoculated in co-culture with the wild type at starting frequencies of up to 50% (Table 1, (Kümmerli et al 2009b, 2015; Kümmerli and Brown 2010).

We first verified that PAO1 produces siderophores in our chronic infection models, and compared production levels with those in iron-limited CAA. Detectable levels of pyoverdine and pyochelin were produced in all environments. Per-cell levels of pyoverdine and pyochelin were (i) positively correlated and (ii) higher in lung and wound infection models than in liquid ASM or SWF (despite lower cells densities in infection models than in corresponding liquid medium) (Figure S1). Consistent with the suggestion of Dumas et al (2013), the pyochelin:pyoverdine ratio was generally higher in chronic infection models and media than in CAA. Replica cultures in *ex vivo* lung and synthetic wounds were more variable than replica cultures in liquid media.

### Social dynamics of siderophore mutants in artificial CF sputum are influenced by genotype and culture time

Mutant and wild-type bacteria were grown in pure culture and each mutant was grown in mixed culture with the wild type, with a starting frequency of 50%, in ASM. After 24 and 48 hours of growth, total population densities were determined and the relative fitness of each mutant in pure and mixed culture calculated. As shown Figure 2a, neither mutant had a relative fitness significantly different from 1 when grown in pure culture. Δ*pvdD* mutant fitness was unaffected by the wild type, but the Δ*pvdD*Δ*pchEF* mutant was outcompeted in mixed culture. These results are not consistent with either mutant acting as a cheat: they do not show a disadvantage when cultured alone, and do not benefit from growth with the wild type.

After 48 hours (Figure 2b), the results were different. The Δ*pvdD* mutant now showed cheating dynamics: it was less fit than the wild type when grown in pure culture but as fit as the wild type in mixed culture. As predicted for a textbook “cheat,” the presence of the Δ*pvdD* mutant reduced total population density: mixed cultures reached similar densities to pure Δ*pvdD* cultures (Figure 2c). The Δ*pvdD*Δ*pchEF* mutant was simply less fit than the wild type regardless of culture condition.

Time course experiments (Figure S2) suggested that the Δ*pvdD* mutant is disadvantaged in pure culture in ASM because it ceases logarithmic growth earlier than the wild type, reaching a lower yield in stationary phase. The Δ*pvdD*Δ*pchEF* mutant has the double disadvantage of a longer lag phase and an earlier cessation of logarithmic growth. This is consistent with the differing fitness of the mutants at 24 vs. 48 hours: continued growth and siderophore production by the wild type from 24-48 h presumably provide an opportunity for the Δ*pvdD* mutant to cheat.

### Siderophore mutants are outcompeted by the wild type in model of CF bronchiolar biofilm

ASM models the chemistry of CF mucus (Palmer et al 2007, Turner et al 2015), but liquid culture lacks realistic spatial structure. To allow bacteria to form biofilm associated with bronchiolar surfaces (Bjarnsholt et al 2013), we repeated the monoculture and 50% co-culture experiments in an *ex vivo* model comprising a section of pig bronchiole cultured in ASM for four days. *P. aeruginosa* forms a loose sleeve of mucoid biofilm around the tissue (Harrison and Diggle in press). The only significant predictor of relative fitness was presence/absence of the wild type: both strains had a relative fitness of 1 in pure culture but showed a trend towards being outcompeted in mixed culture (Figure 3). This trend was significant for the Δ*pvdD*Δ*pchEF* mutant.

### Siderophore mutants do not suffer a long-term fitness disadvantage in synthetic wound fluid

We next repeated the pure/mixed culture experiments in synthetic wound fluid (SWF: Werthén et al 2010). The only significant predictor of relative fitness at 24 hours was presence/absence of the wild type (Figure 4a). The Δ*pvdD* mutant was as fit as the wild type in pure culture, and the Δ*pvdD*Δ*pchEF* mutant slightly and significantly fitter, but both mutants were outcompeted by the wild type in mixed culture. After 48 hours (Figure 4b), there was no difference between genotypes or culture conditions: both mutants had a relative fitness of 1 regardless of wild type presence.

### Siderophore mutants do not have any fitness disadvantage in synthetic chronic wounds

As with the CF lung model, we wished to add spatial structure to SWF to better model a soft tissue infection. SWF was solidified with collagen (Werthén et al 2010) and experiments repeated in the resulting solid plugs.After 48 hours in synthetic wounds, (Figure 5), both mutants had a relative fitness of 1 regardless of culture condition.

## Discussion

Many experiments have been conducted to explore the social evolution of siderophore production by *P. aeruginosa.* These have been used to suggest explanations for the appearance of siderophore mutants in chronic infections; and how this behaviour could be exploited for clinical ends. Our targeted literature review revealed potential restrictions on the generality of predictions made from these experiments about the dynamics of siderophore mutants in nature. First, many experiments used an undefined mutant as a siderophore cheat. We sequenced this strain and found that while it has a deletion of one of the pyoverdine biosynthetic loci, it also carries numerous mutations in genes likely to affect growth and metabolism in unpredictable ways. It is therefore difficult to disentangle the effects of siderophore phenotype vs. other phenotypes on its evolutionary dynamics. Second, most published experiments were carried out in iron-limited minimal broth. Inferences made from these experiments may provide information on the fitness consequences of siderophore loss in acute infections (Granato et al 2016). However, a growing body of evidence strongly suggests that these inferences may not be generalisable to chronic infection contexts, where bacteria experience quite different and idiosyncratic environments that likely affect the cost:benefit ratio of siderophore production, the ability of cells to access each other’s siderophores and the growth rate of bacterial populations. Experiments that dissect the fitness consequences of siderophore loss in environmentally-explicit models of infection could significantly increase our understanding of the natural history of siderophore production, and have enhanced clinical relevance.

We tested whether deletion mutants for pyoverdine or pyoverdine+pyochelin acted as cheats in four laboratory models designed to represent chronic infections: structured and unstructured models of CF lung infections and non-healing soft-tissue wounds. While these models are not perfect, they have been validated with microbiological and chemical data from real infections, and we present them as examples of improved lab models of specific infection contexts.

When increased levels of public goods production increase the population growth rate, or increase carrying capacity, cheating mutants are predicted to be under negative frequency-dependent selection (Ross-Gillespie et al 2007). It is therefore usual to conduct cheating assays using a range of starting frequencies, including <50%. We chose to initially conduct cheating assays using only a 50% starting frequency, as both of the mutants we used have been reported to act as cheats under this condition in iron-limited CAA (Kümmerli et al 2009b, 2015; Kümmerli and Brown 2010). Further, if mutants that cheat from low starting frequencies cannot maintain this advantage as they become more common, then the likely clinical significance of cheating as a determinant of virulence – and the potential power of a “Trojan horse” approach to managing infection – is called into question. Further investigation of different starting frequencies was not necessary: in the case of ASM, the single mutant showed cheating dynamics even at this high starting frequency, and in all other media, both mutants had equal fitness to the wild type in pure culture so cannot be called cheats regardless of the outcome of competition at any starting frequency.

Wild-type *P. aeruginosa* produced siderophores in all of our test environments. However, there was environment-dependent variation in siderophore production, and the ratio of pyochelin:pyoverdine was higher in chronic infection models than in CAA (Figure S1). This is consistent with a suggestion by other authors that pyochelin production may be favoured in chronic infection, where iron is more freely available, or is bound to weaker host chelators than transferrin (Dumas et al 2013, Hunter et al 2013, McCallin et al 2015). The pyoverdine mutant was less fit than the wild type in monoculture in artificial CF sputum after 48 hours’ growth, demonstrating a benefit to pyoverdine production in this environment, and was able to cheat on the wild type. In all other environments tested, there was no effect of losing pyoverdine production on fitness. Production of pyoverdine in these environments may therefore be a maladaptive response, or may have benefits other than simple growth rate enhancement. There was a non-significant trend towards this mutant being outcompeted by the wild type in *ex vivo* bronchiolar biofilm. This may be due to poor biofilm production, even when iron is plentiful (Banin et al 2005, Harrison and Buckling 2009). The pyoverdine+pyochelin mutant was less fit than the wild type in artificial CF sputum and model bronchiolar biofilms, but had no disadvantage in synthetic wound fluid or synthetic wound biofilm.

These results underline the importance of genotype × environment interactions in determining bacterial fitness, and hence the necessity of carefully-designed lab models for predicting the likely consequences of mutations in infection. We previously reported that *P. aeruginosa* QS mutants, which are cheats in an acute infection context (Rumbaugh et al 2009, Wilder et al 2011), are not cheats in an *ex vivo* model of chronic CF lung infection (Harrison et al 2014). Together with the present results, this demonstrates the unreliability of extrapolating predictions about bacterial social evolution from one infection context to another.

So why do siderophore-null mutants arise during chronic infection? Are they cheats, or simply better adapted to local growth conditions? Are they not selected at all, but present transiently and/or at low frequencies? We cannot answer these questions with current data. Careful choice and optimisation of *in vitro* models that allow for long-term evolution experiments in realistic environments will be invaluable in providing answers. Alongside such models there is also a need for (a) quantitative, rather than qualitative, data from patients on the prevalence of siderophore mutants and (b) more considered choice of “wild type” and mutant genotypes, paying particular attention to genotypes that commonly arise during chronic infection. The fitness conseqeunces of siderophore loss could depend on the nature of the mutations involved, and on background genotype. In future, it would be useful to conduct experiments with a range of clinical isolates that are most typical of those seen in patients, and/or with constructed mutants that recapitulate these.

For instance, we and other authors have focussed on experiments with siderophore-null mutants that result from mutations in siderophore biosynthesis. However, a recent study of siderophore mutants isolated from CF patients found that the majority actually carried mutations in the regulatory gene *pvdS* (Andersen et al 2015). Sequencing clones from an *in vitro* evolution experiment also revealed pyoverdine-negative phenotypes that stemmed from *pvdS* mutation (Ross-Gillespie et al 2015, R Kümmerli, personal communication). *PvdS* encodes a sigma factor that positively regulates the expression of the pyoverdine biosynthetic loci and is itself up-regulated by iron starvation (Miyazaki et al 1995), but which also positively regulates the expression of other virulence-related exoproducts (Gaines et al 2007, Hunt et al 2002, Ochsner et al 1996, Wilson et al 2001). Longitudinal sampling from patients (Andersen et al 2015) showed that mutations in siderophore receptors only occur once siderophore-null mutants are present. This was interpreted by the authors as evidence for cheating driving the loss of siderophores. But in the absence of explicit tests for cheating by these isolates in co-culture with co-isolated wild types, in growth media that mimics lung biofilm, the longitudinal pattern presented does not unequivocally prove this conclusion. Other possible explanations include early selection on regulatory loci due to other downstream phenotypes and/or reduced transcriptional costs; a relaxation of purifying selection for functional siderophore receptors in the late stages of infection when iron is plentiful; a bias in the data set towards early isolates which increases the conditional probability that later isolates are also siderophore-null; or a combination of all of these factors.

Our results demonstrate the necessity of evaluating and modelling in-host environments as carefully as possible if we aim to understand in-host microbiology. Lab experiments in simple *in vitro* conditions have limited ecological and clinical validity and so limited predictive power. A plethora of data on the chemical and microbial ecology of chronic infection is now available, providing abundant material for researchers wishing to study the natural history pathogens in the lab (e.g. for CF: Cowley et al 2015, Flynn et al 2016, Folkesson et al 2012, Kyle et al 2015, Quinn et al 2014, Roberts et al 2015, Turner et al 2015, Yang et al 2011). This information is ripe for consideration by microbiologists who – quite justifiably – see great potential for manipulating the in-host ecology of pathogens in order to halt the progression of debilitating chronic infection.

## Acknowledgements

This work was supported by the Universities of Nottingham and Warwick (FH), a Human Frontier Science Program Young Investigators grant to SPD (RGY0081/2012) and a NERC grant to SPD (NE/J007064/1). We thank Pierre Cornelis for the *P. aeruginosa* wild type and siderophore knockout strains, Angus Buckling for PAO9 and Rolf Kümmerli and Stuart West for helpful comments on the manuscript.

## Author contributions

FH & SPD conceived the study and designed experiments. FH, AM and ACS carried out experimental work. FH & AM analysed data. FH drafted the manuscript. All authors contributed to manuscript development.

## Materials and correspondence

Correspondence and requests for materials should be addressed to FH and SPD.

## Supplementary Information

**SuppInfo_1.xls.** Information on the SNPs and deletion found in the genome of PAO6609 (PAO9) (Figure 1). Will be supplied on acceptance or during review process

**SuppInfo_2.xls.** Raw data for Figures 2-5, S1-2. Will be supplied on acceptance or during review process

**Figure S1.**
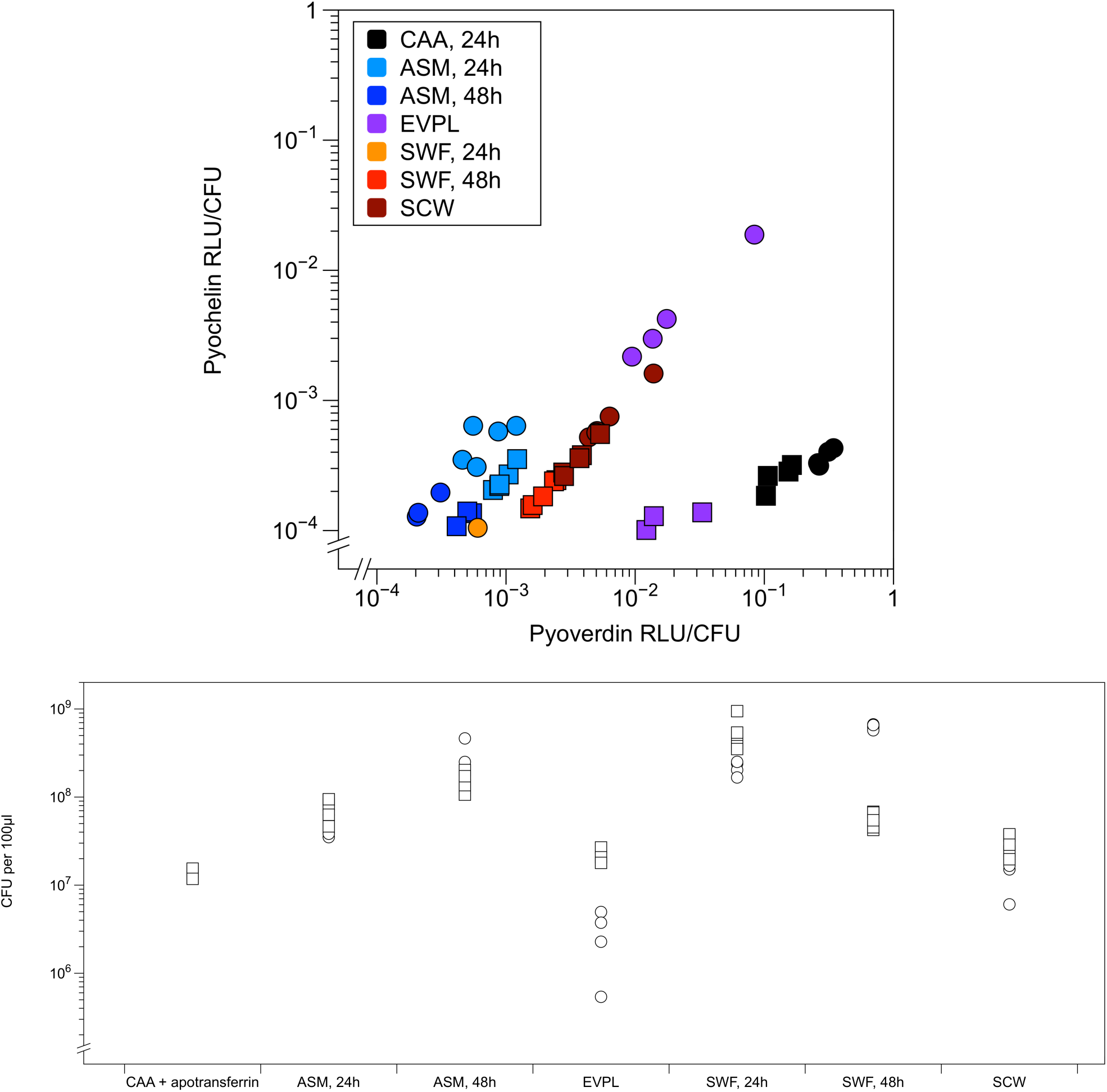
(a) Levels of pyoverdin and pyochelin produced by wild-type *P. aeruginosa* in the different environments explored in this study. Molecules were detected by excitation-emission assays of 100 μl aliquots of culture supernatant and expressed as relative luminescence units (RLU) divided by the number of *P. aeruginosa* colony-forming units (CFU) present in 100 μl of the original culture. Symbols denote experimental block. (b) Bacterial density (CFU per 100 μl) the different environments. Symbols denote experimental block.CAA: casamino acids medium, ASM: artificial sputum medium, EVPL: *ex vivo* pig lung model, SWF: synthetic wound fluid, SCW: synthetic chronic wound model.

**Figure S2.**
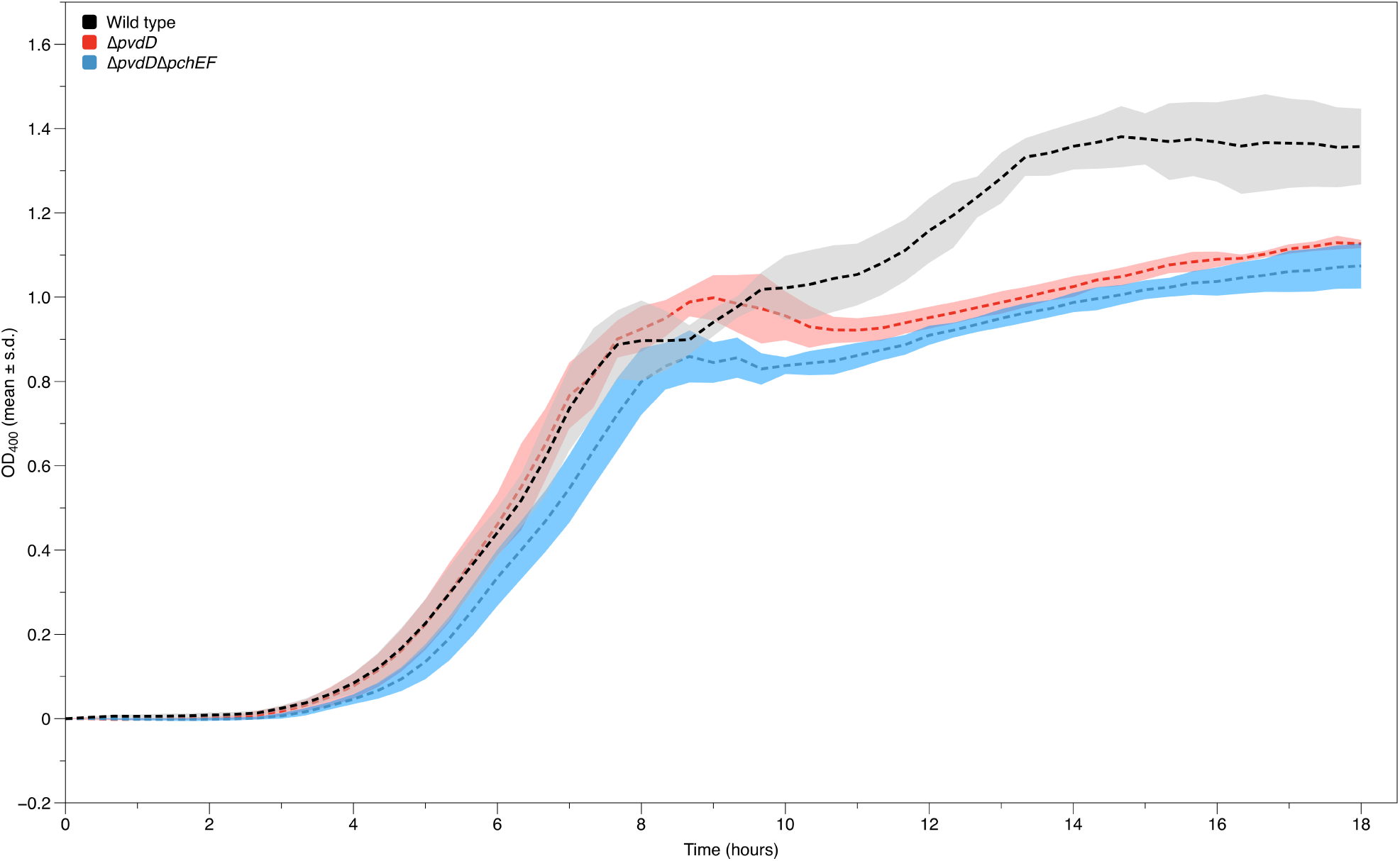
Growth of wild type (black), ∆pvdD (red) and ∆pvdD∆pchEF (blue) P. aeruginosa in 200 μl ASM over 18 hours. Optical density was read at 400 nm to minimise interference from pyoverdine absorbance in wild-type cultures. Lines show means of four replica cultures, shaded areas show ± one standard deviation.

## References

Andersen SB, Marvig RL, Molin S, Krogh Johansen H, Griffin AS (2015). Long-term social dynamics drive loss of function in pathogenic bacteria. Proceedings of the National Academy of Sciences 112: 10756–10761.

Banin E, Vasil ML, Greenberg EP (2005). Iron and *Pseudomonas aeruginosa* biofilm formation. Proceedings of the National Academy of Sciences of the United States of America 102: 11076–11081.

Bjarnsholt T, Alhede M, Alhede M, Eickhardt-Sørensen SR, Moser C, Kühl M et al (2013). The *in vivo* bio-film. Trends in Microbiology 21: 466–474.

Boucher RC (2007). Airway surface dehydration in cystic fibrosis: pathogenesis and therapy. Annual Review of Medicine 58: 157–170.

Brown SP, West SA, Diggle SP, Griffin AS (2009). Social evolution in micro-organisms and a Trojan horse approach to medical intervention strategies. Philosophical Transactions of the Royal Society B: Biological Sciences 364: 3157–3168.

Cowley ES, Kopf SH, LaRiviere A, Ziebis W, Newman DK (2015). Pediatric cystic fibrosis sputum can be chemically dynamic, anoxic, and extremely reduced due to hydrogen sulfide formation. mBio 6 e00767–15.

Croucher NJ, Thomson NR (2010). Studying bacterial transcriptomes using RNA-seq. Current Opinion in Microbiology 13: 619–624.

Darch SE, West SA, Winzer K, Diggle SP (2012). Density-dependent fitness benefits in quorum-sensing bacterial populations. Proceedings of the National Academy of Sciences of the United States of America 109: 8259–8263.

Diggle SP, Griffin AS, Campbell GS, West SA (2007). Cooperation and conflict in quorum-sensing bacterial populations. Nature 450: 411–414.

Dötsch A, Schniederjans M, Khaledi A, Hornischer K, Schulz S, Bielecka A et al (2015). The *Pseudomonas aeruginosa* transcriptional landscape is shaped by environmental heterogeneity and genetic variation. mBio 6: e00749–00715.

Dumas Z, Kümmerli R (2012). Cost of cooperation rules selection for cheats in bacterial metapopulations. Journal of Evolutionary Biology 25: 473–484.

Dumas Z, Ross-Gillespie A, Kümmerli R (2013). Switching between apparently redundant iron-uptake mechanisms benefits bacteria in changeable environments. Proceedings of the Royal Society B: Biological Sciences 280.

Flynn JM, Niccum D, Dunitz JM, Hunter RC (2016). Evidence and role for bacterial mucin degradation in cystic fibrosis airway disease. PLoS Pathogens 12: e1005846.

Folkesson A, Jelsbak L, Yang L, Johansen HK, Ciofu O, Hoiby N et al (2012). Adaptation of *Pseudomonas aeruginosa* to the cystic fibrosis airway: An evolutionary perspective. Nature Reviews Microbiology 10: 841–851.

Foster KR (2005). Hamiltonian medicine: Why the social lives of pathogens matter. Science 308: 1269–1270.

Fox J, Weisberg S (2011). An R Companion to Applied Regression. SAGE Publications.

Frank SA (1998). Foundations of Social Evolution. Princeton University Press. Princeton, NJ.

Friedrich M, Lessnau K-D, Cunha BA (2015). *Pseudomonas aeruginosa* Infections. Medscape Drugs & Diseases. http://emedicine.medscape.com/article/226748.

Gaines JM, Carty NL, Tiburzi F, Davinic M, Visca P, Colmer-Hamood JA et al (2007). Regulation of the *Pseudomonas aeruginosa toxA, regA* and *ptxR* genes by the iron-starvation sigma factor PvdS under reduced levels of oxygen. Microbiology 153: 4219–4233.

Ghoul M, West SA, Diggle SP, Griffin AS (2014). An experimental test of whether cheating is context dependent. Journal of Evolutionary Biology 27: 551–556.

Ghoul M, West SA, McCorkell FA, Lee Z-B, Bruce JB, Griffin AS (2016). Pyoverdin cheats fail to invade bacterial populations in stationary phase. Journal of Evolutionary Biology 29: 1728–1736.

Ghysels B, Dieu BTM, Beatson SA, Pirnay JP, Ochsner UA, Vasil ML et al (2004). FpvB, an alternative type I ferripyoverdine receptor of *Pseudomonas aeruginosa*. Microbiology 150: 1671–1680.

Goodman AL, Kulasekara B, Rietsch A, Boyd D, Smith RS, Lory S (2004). A signaling network reciprocally regulates genes associated with acute infection and chronic persistence in *Pseudomonas aeruginosa*. Developmental Cell 7: 745–754.

Granato ET, Harrison F, Kümmerli R, Ross-Gillespie A (2016). When is a bacterial “virulence factor” really virulent? bioRxiv. doi: 10.1101/061317

Griffin AS, West SA, Buckling A (2004). Cooperation and competition in pathogenic bacteria. Nature 430: 1024–1027.

Hamilton WD (1964). The genetical evolution of social behaviour I & II. Journal of Theoretical Biology 7: 1–52.

Harrison F, Browning LE, Vos M, Buckling A (2006). Cooperation and virulence in acute *Pseudomonas aeruginosa* infections. BMC Biology 4: 21.

Harrison F, Paul J, Massey RC, Buckling A (2008). Interspecific competition and siderophore-mediated cooperation in *Pseudomonas aeruginosa*. ISME Journal 2: 49–55.

Harrison F, Buckling A (2009). Siderophore production and biofilm formation as linked social traits. ISME Journal 3: 632–634.

Harrison F, Buckling A (2011). Wider access to genotypic space facilitates loss of cooperation in a bacterial mutator. PLoS ONE 6: e17254.

Harrison F, Muruli A, Higgins S, Diggle SP (2014). Development of an *ex vivo* porcine lung model for studying growth Virulence, And signaling of *Pseudomonas aeruginosa*. Infection and Immunity 82: 3312–3323.

Harrison F, Diggle S (in press). An *ex vivo* lung model to study bronchioles infected with *Pseudomonas aeruginosa* biofilms. Microbiology. doi: 10.1099/mic.0.000352.

Hirsch EB, Tam VH (2010). Impact of multidrug-resistant Pseudomonas aeruginosa infection on patient outcomes. Expert review of pharmacoeconomics & outcomes research 10: 441–451.

Hohnadel D, Haas D, Meyer JM (1986). Mapping of mutations affecting pyoverdine production in *Pseudomonas aeruginosa*. FEMS Microbiology Letters 36: 195–199.

Hothorn T, Bretz F, Westfall P (2008). Simultaneous inference in general parametric models. Biometrical Journal 50: 346–363.

Hunt TA, Peng W-T, Loubens I, Storey DG (2002). The *Pseudomonas aeruginosa* alternative sigma factor PvdS controls exotoxin A expression and is expressed in lung infections associated with cystic fibrosis. Microbiology 148:3183–3193.

Hunter RC, Asfour F, Dingemans J, Osuna BL, Samad T, Malfroot A, Cornelis P, Newman, DK (2013). Ferrous iron is a significant component of bioavailable iron in cystic fibrosis airways. mBio 4: e00557–13.

Jiricny N, Diggle SP, West SA, Evans BA, Ballantyne G, Ross-Gillespie A et al (2010). Fitness correlates with the extent of cheating in a bacterium. Journal of Evolutionary Biology 23: 738–747.

Kümmerli R, Gardner A, West SA, Griffin AS (2009a). Limited dispersal, budding dispersal, and cooperation: an experimental study. Evolution 63: 939–949.

Kümmerli R, Jiricny N, Clarke LS, West SA, Griffin AS (2009b). Phenotypic plasticity of a cooperative behaviour in bacteria. Journal of Evolutionary Biology 22: 589–598.

Kümmerli R, Griffin AS, West SA, Buckling A, Harrison F (2009c). Viscous medium promotes cooperation in the pathogenic bacterium *Pseudomonas aeruginosa*. Proceedings of the Royal Society B: Biological Sciences 276: 3531–3538.

Kümmerli R, Brown SP (2010). Molecular and regulatory properties of a public good shape the evolution of cooperation. Proceedings of the National Academy of Sciences of the United States of America 107: 18921–18926.

Kümmerli R, Van Den Berg P, Griffin AS, West SA, Gardner A (2010). Repression of competition favours cooperation: experimental evidence from bacteria. Journal of Evolutionary Biology 23: 699–706.

Kümmerli R, Ross-Gillespie A (2014). Explaining the sociobiology of pyoverdin producing *Pseudomonas:* a comment on Zhang and Rainey (2013). Evolution 68: 3337–3343.

Kümmerli R, Santorelli LA, Granato ET, Dumas Z, Dobay A, Griffin AS et al (2015). Co-evolutionary dynamics between public good producers and cheats in the bacterium *Pseudomonas aeruginosa*. Journal of Evolutionary Biology 28: 2264–2274.

McCallin K, Cowley E, Reyes MC, Van Sambeek L, Hunter R, Asfour F et al (2015). Sputum iron levels during cystic fibrosis pulmonary exacerbation: a longitudinal study. American Thoracic Society International Conference Abstracts B52: Pediatric Cystic Fibrosis pp A3343–A3343.

Lenth RV (2013). lsmeans: R Package Version 1.06-05.

Miyazaki H, Kato H, Nakazawa T, Tsuda M (1995). A positive regulatory gene, *pvdS*, for expression of pyoverdin biosynthetic genes in *Pseudomonas aeruginosa* PAO. Molecular and General Genetics MGG 248: 17–24.

Mund A, Diggle SP, Harrison F (2016). The fitness of *Pseudomonas aeruginosa* quorum sensing signal cheats is influenced by the diffusivity of the environment. bioRxiv. doi: 10.1101/082230

Ochsner UA, Johnson Z, Lamont IL, Cunliffe HE, Vasil ML (1996). Exotoxin A production in *Pseudomonas aeruginosa* requires the iron-regulated *pvdS* gene encoding an alternative sigma factor. Molecular Microbiology 21: 1019–1028.

Palmer KL, Mashburn LM, Singh PK, Whiteley M (2005). Cystic fibrosis sputum supports growth and cues key aspects of *Pseudomonas aeruginosa* physiology. Journal of Bacteriology 187: 5267–5277.

Palmer KL, Aye LM, Whiteley M (2007). Nutritional cues control *Pseudomonas aeruginosa* multicellular behavior in cystic fibrosis sputum. Journal of Bacteriology 189: 8079–8087.

Quinn RA, Lim YW, Maughan H, Conrad D, Rohwer F, Whiteson KL (2014). Biogeochemical forces shape the composition and physiology of polymicrobial communities in the cystic fibrosis lung. mBio 5: e00956–13.

Raymond B, West SA, Griffin AS, Bonsall MB (2012). The dynamics of cooperative bacterial virulence in the field. Science 336: 85–88.

Rella M, Mercenier A, Haas D (1985). Transposon insertion mutagenesis of *Pseudomonas aeruginosa* with a Tn5 derivative: application to physical mapping of the *arc* gene cluster. Gene 33: 293–303.

Roberts AEL, Kragh KN, Bjarnsholt T, Diggle SP (2015). The limitations of *in vitro* experimentation in understanding biofilms and chronic infection. Journal of Molecular Biology 427: 3646–3661.

Ross-Gillespie A, Gardner A, West SA, Griffin AS (2007). Frequency dependence and cooperation: Theory and a test with bacteria. American Naturalist 170: 331–342.

Ross-Gillespie A, Gardner A, Buckling A, West SA, Griffin AS (2009). Density dependence and cooperation: theory and a test with bacteria. Evolution 63: 2315–2325.

Ross-Gillespie A, Dumas Z, Kümmerli R (2015). Evolutionary dynamics of interlinked public goods traits: an experimental study of siderophore production in *Pseudomonas aeruginosa*. Journal of Evolutionary Biology 28: 29–39.

Rumbaugh KP, Diggle SP, Watters CM, Ross-Gillespie A, Griffin AS, West SA (2009). Quorum sensing and the social evolution of bacterial virulence. Current Biology 19: 341–345.

Stover CK, Pham XQ, Erwin AL, Mizoguchi SD, Warrener P, Hickey MJ et al (2000). Complete genome sequence of *Pseudomonas aeruginosa* PAO1, an opportunistic pathogen. Nature 406: 959–964.

Team RDC (2016). R: A Language and Environment for Statistical Computing. R Foundation for Statistical Computing, Vienna, Austria. http://www.R-project.org.

Turner KH, Wessel AK, Palmer GC, Murray JL, Whiteley M (2015). Essential genome of *Pseudomonas aeruginosa* in cystic fibrosis sputum. Proc Natl Acad Sci U S A 112: 4110–4115.

Tyrrell J, Callaghan M (2016). Iron acquisition in the cystic fibrosis lung and potential for novel therapeutic strategies. Microbiology 162: 191–205.

Van Opijnen T, Bodi KL, Camilli A (2009). Tn-seq: high-throughput parallel sequencing for fitness and genetic interaction studies in microorganisms. Nat Methods 6.

Werthén M, Henriksson L, Jensen PØ, Sternberg C, Givskov M, Bjarnsholt T (2010). An *in vitro* model of bacterial infections in wounds and other soft tissues. APMIS 118: 156–164.

West SA, Diggle SP, Buckling A, Gardner A, Griffin AS (2007). The social lives of microbes. Annual Review of Ecology, Evolution, and Systematics. pp 53–77.

Whiteley M, Bangera MG, Bumgarner RE, Parsek MR, Teitzel GM, Lory S et al (2001). Gene expression in *Pseudomonas aeruginosa* biofilms. Nature 413: 860–864.

Wilder CN, Diggle SP, Schuster M (2011). Cooperation and cheating in *Pseudomonas aeruginosa:* the roles of the las, rhl and pqs quorum-sensing systems. ISME Journal 5: 1332–1343.

Wilson MJ, McMorran BJ, Lamont IL (2001). Analysis of promoters recognized by PvdS, an extracytoplas-mic-function sigma factor protein from *Pseudomonas aeruginosa*. Journal of Bacteriology 183: 2151–2155.

Yang L, Jelsbak L, Marvig RL, Damkiær S, Workman CT, Rau MH et al (2011). Evolutionary dynamics of bacteria in a human host environment. Proceedings of the National Academy of Sciences of the United States of America 108: 7481–7486.

Zhang X-X, Rainey PB (2013). Explaining the sociobiology of producing *Pseudomonas*. Evolution 67: 3161–3174.

